# Hunter-gatherer foraging networks promote information transmission

**DOI:** 10.1101/2021.04.29.442031

**Authors:** Ketika Garg, Cecilia Padilla-Iglesias, Nicolás Restrepo Ochoa, V. Bleu Knight

**Author notes:** Contributed equally (Ketika Garg, Cecilia Padilla-Iglesias).

## Abstract

Central-place foraging, where foragers return to a central location (or home), is a key feature of hunter-gatherer social organization. Central-place foraging could have significantly changed hunter-gatherers’ use of space and mobility, and altered social networks and increased opportunities for information exchange. We evaluated whether central-place foraging patterns facilitate information transmission and considered the potential roles of environmental conditions and mobility strategies. We built an agent-based central-place foraging model where agents move according to a simple optimal foraging rule, and can encounter other agents as they move across the environment. They either forage close to their home within a given radius or move the location of their home to new areas. We analyzed the interaction networks arising across different environments and mobility strategies. We found that, at intermediate levels of environmental heterogeneity and mobility, central-place foraging increased global and local network efficiencies as well as the rate of contagion-based information transmission (simple and complex). We also assessed the effect of population density on the resultant networks and found that central-place mobility strategies can further improve information transmission in larger populations. Our findings suggest that the combination of foraging and movement strategies, as well as the underlying environmental conditions that characterized early human societies, may have been a crucial precursor in our species’ unique capacity to innovate, accumulate and rely on complex culture.

## 1 Introduction

One of the pivotal transitions in human evolution is our ability to innovate, accumulate and rely on complex, cumulative culture (1; 2; 3; 4). Recent evidence from hunter-gatherer societies (5; 6) has suggested that changes in our ancestors’ social networks and connectivity could have promoted such a transition by facilitating an efficient exchange and transmission of cultural information. Given that the frequency and nature of social interactions between hunter-gatherers would have been affected by their movement and spatial distribution patterns, researchers have proposed that divergences in foraging behavior, coupled with ecological changes, could have led to changes in the dynamics of social interactions and hence patterns of social organization (7; 8; 9). However, the precise impact of hunter-gatherer foraging and movement behavior on emergent social networks and their ability to transmit information is still not thoroughly understood (but see 10; 11).

Central-place foraging marks a critical behavioral change between the foraging styles of early hominins and our closest great ape relatives (12; 13; 14) that would have modified their movement and consequently spatial and social patterning.

Non-human Great Apes (henceforth Great Apes) tend to consume food when they find it (‘point-to-point’ foraging), make sleeping nests at variable locations and have shorter foraging trips (15). On the other hand, hunter-gatherers establish residential camps or central places around which they systematically forage and bring the food they collect during foraging trips (or logistical forays) back to their camps to share and process it with camp members (‘central-place’ foraging) (16; 17). In addition, human foragers can make longer foraging trips and periodically move the location of their residential camps to access new resource areas with little overlap in the foraging-radii between successive camps. These properties result in an expansion of their overall home-ranges during their lifetimes (15) compared to other primates who spend most of their adult lives within the same area, leading to a more restricted use of space (18; 19; 20; 21; 22).

Such differences in mobility could have altered spatial patterns and dynamics of social interactions and led to more complex social structures. In particular, we hypothesize that central-place foraging could have played an essential role in the subsequent development of multi-level sociality. In multi-level organizations, sets of multiple core units (like families) repeatedly coalesce, intermix and disperse, giving rise to relatively fluid local bands that are embedded in higher-level interconnected regional networks(23; 24; 25; 26; 27; 28). These extended, flexible, and fluid social landscapes would have increased the likelihood of interactions, social learning, and information exchange compared to the rest of the Great Apes (2; 29).

However, hunter-gatherer foraging and mobility decisions (e.g. daily trips, residential movements) are influenced by their resource environments as well as various costs (e.g., traveling costs) and benefits (e.g., resource abundance) of their foraging activities (30; 31). Hunter-gatherer bands may be able to afford greater sedentism in rich environments where there are plenty of resources available within their foraging-radii (or home-ranges) (32). In contrast, unproductive may require bands to move their camps multiple times a year due to resource depletion within their foraging radii (33). In addition, if resources are homogeneously distributed, ethnographic studies have shown that bands tend to predominantly rely on short and frequent residential moves (17). In these settings, the frequency of encounter with other bands and thus, the inter-connectivity between them could decrease. Conversely, if resources are heterogeneously distributed and some areas are more resource-rich than others, bands may aggregate in key locations from which they conduct daily trips to procure resources (30), potentially generating more opportunities for interactions (34).

In this paper, we model and compare point-to-point and central-place foraging (with different home-range radii) behavior across a range of environments. We investigate the effect of central-place foraging and mobility on the interaction patterns between foraging units and the subsequent social networks that are formed due to foraging units coinciding on resources. We then test the efficiency of information transmission in the networks that emerge from the different mobility regimes and environments. Previous theoretical and computational models have explored the effects of environmental heterogeneity on social networks emerging from foraging behavior across different environments (35) and hunter-gatherer mobility on cultural transmission (10; 36). However, models explicitly linking foraging strategies, environmental features, and hunter-gatherer interaction networks remain lacking. Our work illustrates a direct connection between environmental conditions, foraging behavior, and information flow in hunter-gatherer social networks, thereby providing insights into the evolutionary origins of our species’ unique ability to innovate, accumulate and rely on complex culture.

## 2 Methods

### 2.1 Model Description

We investigated how central-place-foraging behavior would affect the emergent interaction networks across environments. Previous work by Ramos-Fernández et al. (35) modeled the effect of environmental heterogeneity on the interaction networks that emerge from multiple agents foraging independently (representing spider monkeys). The authors showed that a complex social structure with fission-fusion properties, resembling those observed in field studies among real spider monkey societies, could emerge simply from optimal foraging rules in heterogeneous environments.

Our model (henceforth central-place model), like the model from Ramos-Fernández et al. (35) (henceforth point-to-point model), is executed in a two-dimensional environment that ranges from 0 to 1, and comprises 50,000 uniformly distributed patches. Each patch was initially assigned resource content, *k*_*i*_ ≥ 1, drawn from a normalized power-law probability distribution, *P* (*k*) ≈ *Ck*^*−β*^ where the exponent *β* determines the distribution of resource content and the total resource abundance (see Supplementary Methods section and Fig. 1) and 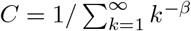 is the normalization constant. Following this equation, the richness of an environment (or abundance) and its heterogeneity (or distribution) co-vary and are determined by *β*.

**Figure 1:**
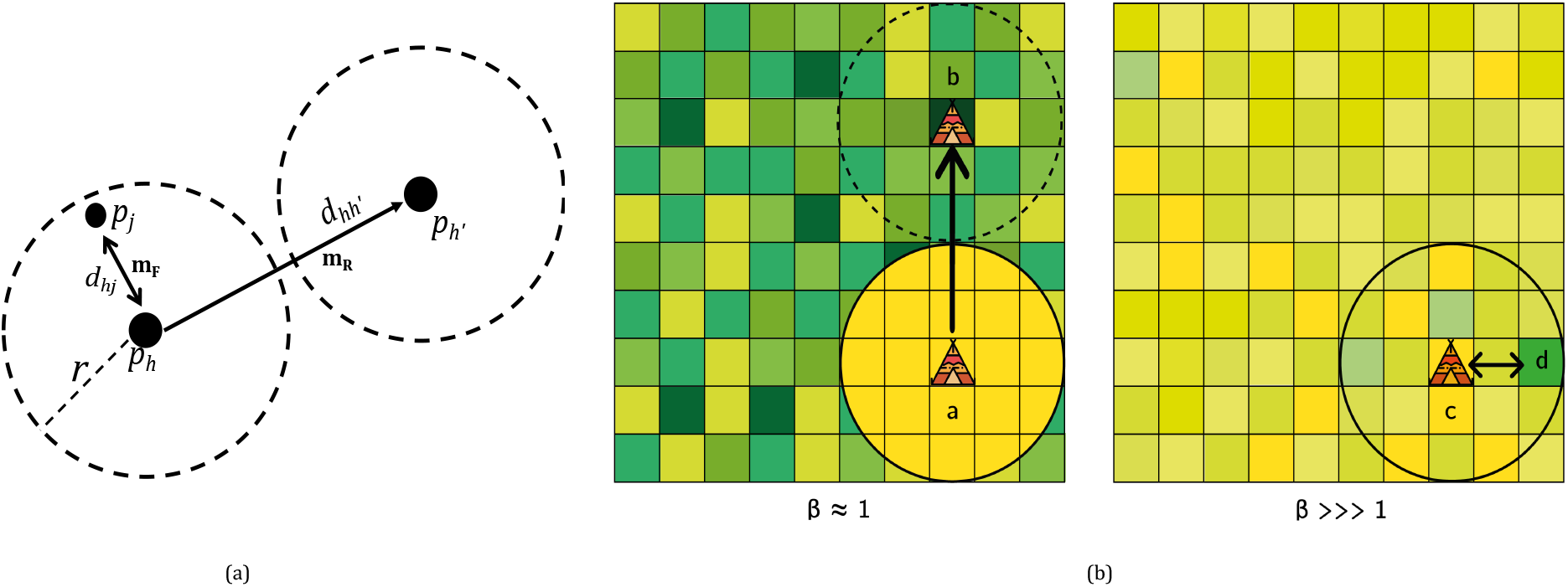
Model Description. ***(a)***: A schematic representation of agent movement in the model. Agents can make foraging movements (*m*_*F*_) within a radius *r* from their home (*p*_*h*_) to a new patch (*p*_*j*_) or residential movements (*m*_*R*_) to a new home (*p*_*h*′_). ***(b)***: An illustration of the variations in resource environments modulated by the parameter *β*. A low value of *β* results in a rich (dark-green patches) and heterogeneous environment (left), whereas a very high value of *β* results in a scarce (yellow patches) and homogeneous environment (right). When food depletes within an agent’s radius (yellow patches), it moves its residence (*a* → *b*). Otherwise, it continues to forage within its radius (*c* ↔ *d*).

When *β* ≈ 1, *k* has a broad range with high values of *k*, patches vary widely in their resource content, and the environment is abundant with many rich patches. Conversely, *β* ≫ 1 corresponds to smaller values and a restricted range of *k* that result in an environment composed of scarcer resources that are homogeneously distributed across patches. Patches deplete by a unit every time-step a foraging-unit spends in it, and the patches do not regenerate (see Fig S5 for more information on resource depletion).

Each agent in our simulation represents a monogamous, nuclear family/foraging unit (adult male, female and dependent offspring) which are the core, indivisible units of social organization across hunter-gatherer societies (37). Since ties between individuals from different families would result in a tie between the families, agents are assumed to forage and move as a single foraging-unit. The model is initialized with agents independently and randomly distributed across the patches. Foraging-units followed a rule whereby they move to a new patch (*p*_*j*_) from a depleted patch (*p*_*i*_) such that it minimized the cost/gain ratio (*d*_*ij*_*/k*_*j*_), where *d*_*ij*_ is the distance between the patches and *k*_*j*_ is the resource content of *p*_*j*_. Our model (Fig 1a) modified this resource-maximization rule to implement central-place foraging and distinguished between foraging (or logistical) and residential moves (16; 17).

In our model, foraging-units move to fixed home locations from which they exploit the surrounding local environment in their foraging radius before moving to another home location. Every foraging-unit had complete knowledge of resources, a randomly allocated home location (i.e. central place), and a foraging area with a given radius, *r*. Foraging-units could forage and change their home location based on the following rules: When foraging-units were on a patch with no food left, they made foraging moves (*m*_*F*_) to a patch (*p*_*j*_) within *r* such that the cost/gain ratio (*d*_*hj*_*/k*_*j*_) was minimized, where *d*_*hj*_ is distance from the current home (*p*_*h*_), and *k*_*j*_ is the resource content of *p*_*j*_ (Fig 1b (right)). Before every move, foraging-units compared the cost/gain ratio of patches outside the radius to patches within the radius. When the resource quality within *r* diminished compared to the rest of the environment (Fig 1b (left)), instead of making a foraging move to profitable patches outside their radius, foraging-units made a residential move. Residential moves (*m*_*R*_) allowed foraging-units to select a new home (*p*_*h*_) that minimized (*d*_*hh*′_ */k*_*h*′_) but was far enough from the current base (*d*_*hh*′_ ≥ 2 * *r*) to avoid overlap (10). Each time-step that a foraging-unit coincided with another foraging-unit on a patch, they formed a social network tie or added a unit of weight to an existing tie.

To assess how the combination of environmental heterogeneity and central-place foraging strategies affect the emergent social networks, we varied the resource exponent, *β* parameter to take values between 1.5 and 4.5 the foraging radius, *r* to assume values of 1, 0.1, 0.01, and 0.001. Furthermore, we tested the effect of population size by running the model with 50, 100, and 200 foraging-units (Fig. S1). Whilst our main motivation with this manipulation was to explore the effect of changing population sizes on our model results, the values we chose are ethnographically meaningful. We mostly focused on population sizes of 100 foraging units/families (that correspond to 500 individuals) which have been documented widely (38) and are assumed to represent the average size of hunter-gatherer regional bands or groups (18). Populations of size 200 correspond to some estimates of the size of entire ethnic populations (or metapopulations) (as compiled in Lehmann et al. (39)). Finally, populations of 50 families (200-250 individuals) represent of a lower limit for hunter-gatherer populations to remain viable (40).

We ran 50 simulations for each parameter combination and the point-to-point model and extracted the weighted social networks formed throughout the simulation as well as at the end of the simulation. To ensure that our results are not an artifact of the chosen time-steps, we conducted sensitivity analyses, running each parameter combination for 1000 time-steps (available in the supplementary materials). We found the results to be consistent over longer time-steps and thus, report the results from the first 100 time-steps in the following text.

### 2.2 Networks

We extracted the final networks formed from the sum of all interactions by the end of each run. We provide complete summary statistics of each parameter combination’s resulting networks in the supplementary materials (Tables S4-S5). As a robustness check, we also look at how the networks develop across the simulations. For 100 time-steps, we examine the networks after each interval of 10 time-steps. For 1000 time-steps, we increase that window to a 100 time-steps.

#### 2.2.1 Efficiency

We tested the networks for their ability to transmit information by measuring their global and local efficiencies (41). The efficiency measures have been used across various studies to investigate the transmission of social and cultural information in various networks, including hunter-gatherer social networks (5). Global efficiency indicates a network’s ability to transmit information across the entire network and is inversely related to the characteristic path length (or the average distance between nodes). Latora and Marchiore (41) define a graph’s global efficiency as the inverse of the sum of the shortest paths between all nodes i and j:

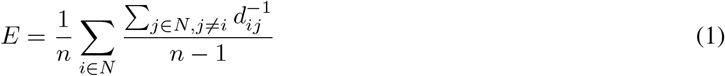

Where *N* is the set of all nodes in the network, *n* is the total number of nodes, and *d* is the shortest distance between two nodes. On the other hand, local efficiency relates to the clustering coefficient of a network (i.e., the degree to which a node’s local neighborhood is inter-connected). It measures the average global efficiency of subgraphs and denotes how well each local neighborhood can exchange information within itself. We modified the efficiency measures to incorporate weights (for a more detailed description, see SM).

#### 2.2.2 Contagion simulations

We calculated the proportion of agents in the extracted network at the end of simulations that acquired the diffusing information after 5000 time-steps through simple and complex contagion. Simple contagion models a perfect transmission of information independent of the number of novel interactions, where a single interacting event is sufficient for information transmission. Hence, an agent’s probability of acquiring information is proportional to the number of its neighbors with the information and the strength of their connections. Accounting for edge weights, the probability of acquiring the information for individual *i*, in any given turn, is:

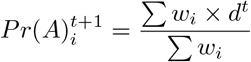

Where *w*_*i*_ is a vector of the edge weights that *i* shares with its neighbors and *d* is a vector of same length containing 1 if the corresponding neighbor has acquired the information or 0 otherwise at time, *t*.

Complex contagion, in contrast, represents a mode of transmission that is more well-suited to capture the diffusion of costly or difficult social behaviors that need reiterated affirmation (42; 43). Here, the probability of acquisition now rises exponentially as more neighbors acquire the information,

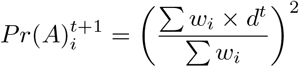

## 3 Results

### 3.1 Environmental factors affect the efficiency of information transmission in networks

In line with the results from Ramos-Fernández et al. (35), we found that environmental heterogeneity strongly influenced the networks formed, with *β* = 2.5 generating the most efficient networks (*Ē*_*global*_ = 0.13, *Ē*_*local*_ = 0.65). In environments of (*β* ≈ 1) where many rich resource patches were available, foraging-units had very low mobility (see next section for mobility results) and stayed fixed at a rich resource patch for long durations. In the homogeneous environment of *β* = 4.5, every patch had low resource value, and foraging-units depleted patches quickly. They frequently moved across the environment resulting in low interaction rates (as evidenced by density of connections) with other foraging-units. However, at intermediate heterogeneity and resource abundance (*β* = 2.5), foraging-units coincided at many different rich patches available in the environment and formed stronger social ties. This can be evidenced by the high number of total interactions between foraging-units per time-step (Fig. S4) that increased the network’s local efficiency. On the other hand, intermediate number of rich patches also enabled more movement and unique interactions between the foraging-units(Fig. S12-13) that made the network more expansive and increased its global efficiency. Increasing the population’s size further increased the rate of interactions between the foraging-units and thus, the network efficiencies (Fig. S1).

### 3.2 Central-place foraging increases global and local network efficiency

We found that point-to-point foraging created networks comprised of isolated foraging-units with very high local efficiency (or clustering) but low global efficiency. These networks contained strongly connected small sub-groups of foraging-units that were distributed across the environment with few to no connections between them. In contrast, central-place foraging increased the number of unique interactions between foraging-units (Fig S12-13) and created ties between otherwise unconnected sub-groups (or components) that resulted in a more connected and expansive network (Figure S7). A completely disconnected graph has a number of components equal to its nodes, while a fully connected graph has a single component. We found that the ties between sub-groups decreased the number of components and increased networks’ global efficiency while maintaining high local efficiency. On the one hand, this formed strongly bonded local groups, and on the other hand, large-scale, interconnected regional networks such as the ones observed among ethnographic hunter-gatherers (26; 2; 44; 5)

To explore the effect of different radii of central-place foraging, we compared the different mobility regimes (frequency and magnitude of residential and foraging moves) across environments and radii 4 (Tables S2-S3).

When the foraging radius was small (*r* = 0.001), at intermediate levels of environmental heterogeneity (*β* = 2.5), the observed foraging pattern closely resembled point-to-point foraging (15), where foraging-units don’t return to a central-place, make short residential moves 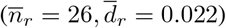 and fewer and shorter foraging moves 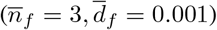 The resulting networks from *r* = 0.001 comprised many (≈ 24) densely connected sub-groups of high local efficiencies (*Ē*_*local*_ = 0.73)(Table S4). Nonetheless, these dense sub-groups lacked connections between them with a maximum of 3 sub-groups connected to each other (Table S5).

Increasing the foraging radius to intermediate values (*r* = 0.01 and *r* = 0.1) resulted in foraging-units making longer, and more frequent foraging moves combined with longer but fewer residential moves. At *r* = 0.01, a small increase in residential mobility 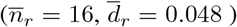 created a few longer connections between the dense sub-groups. These connections resulted in more sub-groups being connected (4−5) and a more expansive network that increased the global efficiency (*Ē*_*global*_ = 0.11). However, the network still remained highly cliquish with high local efficiency (*Ē*_*local*_ = 0.75).

As the foraging radius and use of space further increased (*r* = 0.1), foraging-units moved less frequently 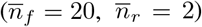 but undertook longer moves 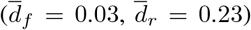. This change in mobility increased the long-range connections between fewer (≈ 15) but larger and interconnected sub-groups (6 − 8). The resultant sub-group structure made the network substantially more efficient at the global scale (*Ē*_*global*_ = 0.24) while maintaining considerable local efficiencies (*Ē* _*local*_ = 0.50)(Tables S4-S5). However, we found that both efficiencies decreased compared to intermediate radii (*Ē*_*local*_ = 0.44, *Ē*_*global*_ = 0.12) when the foraging-units had a very large foraging radius (*r* = 1). In the absence of residential moves, the foraging-units remained tethered to their original home and traversed longer foraging moves 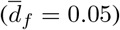 to find food. The longer moves helped create long-range connections between foraging-units that resulted in a large number of connected sub-groups (7 − 11) and a more globally efficient network than the point-to-point model (*Ē*_*global*_ = 0.07). But the strong tethering decreased the overall use of space and the probability of coinciding with others for longer durations, resulting in fewer and weaker connections between sub-groups (see Fig. S2-S3) with low local efficiencies.

In environments where the habitat quality was lower and patches were more homogeneous in their resource content, foraging-units coincided on patches less frequently and for a shorter amount of time that resulted in fewer interactions (Fig S6, S12-13). At *β* = 3.5 when fewer patches were rich, foraging mobility increased with many shorter moves within foraging-radii for all radii, while residential mobility increased with longer moves 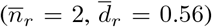 for *r* = 0.1, but decreased for smaller radii with shorter and similar number of moves 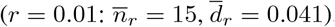, or shorter and more frequent moves 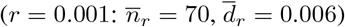 This effect led to a decrease in both global and local efficiencies across radii (*Ē*_*global*_ = 0.04, *Ē*_*local*_ = 0.3) from *β* = 2.5. However, for radius *r* = 0.01, the decrease in the local efficiencies was lesser when compared to the other radii. For *r* = 0.01, foraging-units moved within a space that was small enough to increase the rate of interactions but large enough to find rich patches. When the radius increased (*r* = 0.1, 1) or decreased (*r* = 0.001), foraging-units either traveled longer distances and were dispersed in a larger area or were too restricted in their space use to find enough food and continually changed their residence.

Taken together, these data illustrate that central-place foraging-units were restricted in their movements that led to strongly connected sub-groups. However, the longer residential moves allowed connections to form between the sub-groups to varying extents that were missing in the point-to-point model. We found that intermediate levels of overall mobility 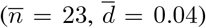 with few long moves, and many shorter moves, for example in *β* = 2.5 and *r* = 0.1, created networks that were efficient at both global and local scales. As the frequency of overall movement decreased with longer moves 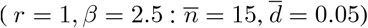 or increased with shorter moves 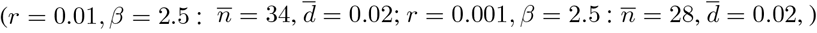, networks lacked dense, long-range connections necessary for global efficiency with either highly locally efficient but fragmented networks or sparsely connected sub-groups. Finally, when rate of mobility was very low (highly frequent but very short moves, or rare and and short moves), for example in *β* = 1.5 and *β* = 4.5 (all radii), foraging-units rarely interacted with each other, and both global and local efficiencies tended to 0. Altogether, based on our results, we can predict that central-place foraging with an intermediate radius/mobility regime (between 0.01 - 0.1) should maximize both efficiencies (Fig 2 inset). Furthermore, our sensitivity analyses indicate that this result is robust over longer time-steps (see Supplementary Results for more information).

**Figure 2:**
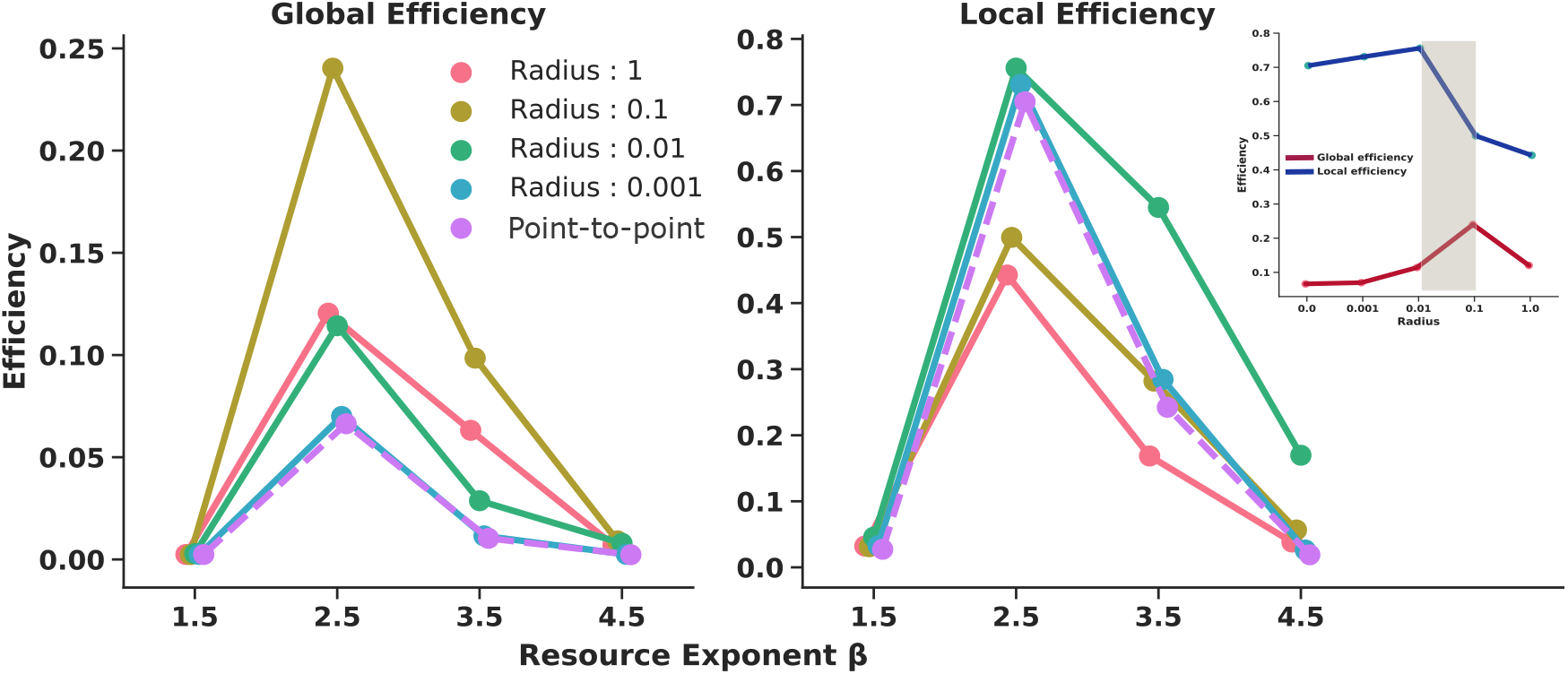
Network Efficiencies after 100 time-steps. The plot shows the average global (left) and local (right) efficiencies of the networks as a function of environmental heterogeneity for each radius. The inset shows the relationship between efficiencies and radius for *β* = 2.5. Shaded region corresponds to intermediate radii that balances global and local efficiencies.

### 3.3 Population size and mobility affect network efficiency

We also tested the effect of varying population sizes on the resultant networks by simulating populations of 50, 100, 200 foraging-units. We found that as population size increased, regardless of the radius, the local efficiencies of the networks also increased (Fig.5a). An increase in population size led to a higher rate of coincidence between foraging-units on patches that created denser connections. This effect was stronger when the radius was smaller (*r* ≤ 0.01) because the agents were restricted within smaller areas that led to repetitive interactions and added weight to local connections.

**Figure 3:**
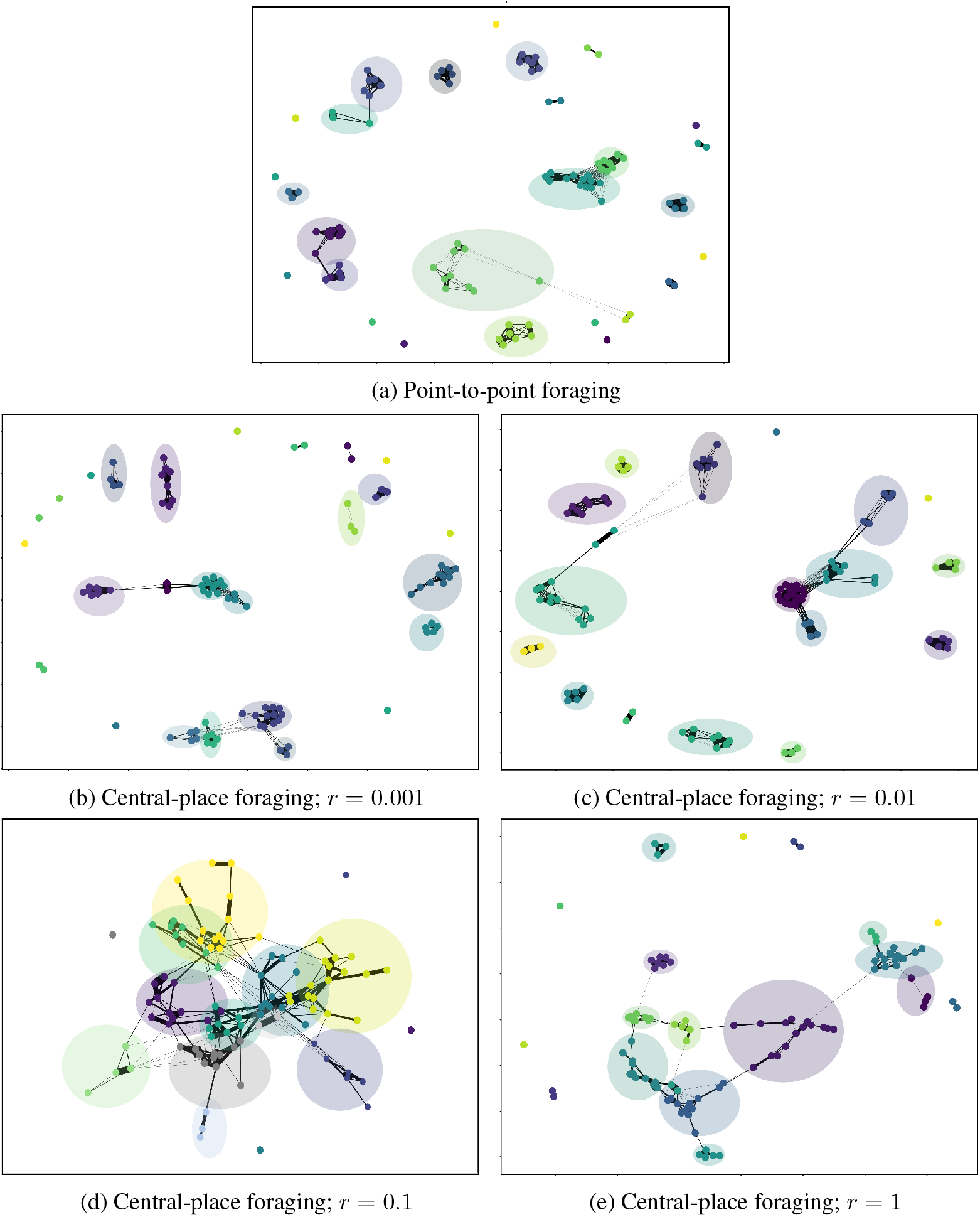
Emergent Networks from 100 foraging-units after 100 time-steps. The plot shows an example of weighted networks that emerge from different foraging behaviors in *β* = 2.5 environment (clockwise: *Point-to-point, r* = 0.001, *r* = 0.01, *r* = 0.1, *r* = 1). Node colors depict the different sub-groups detected by Louvain community-detection method (SI Text). Different communities and the overlap between them are also shown by circles around each community. Edge widths depict the edge weights, with thicker edges representing stronger bonds, and finer edges representing weaker bonds. Distance between nodes also depict the strength of connections. Networks were made using spring layout from *NetworkX* package in Python.

**Figure 4:**
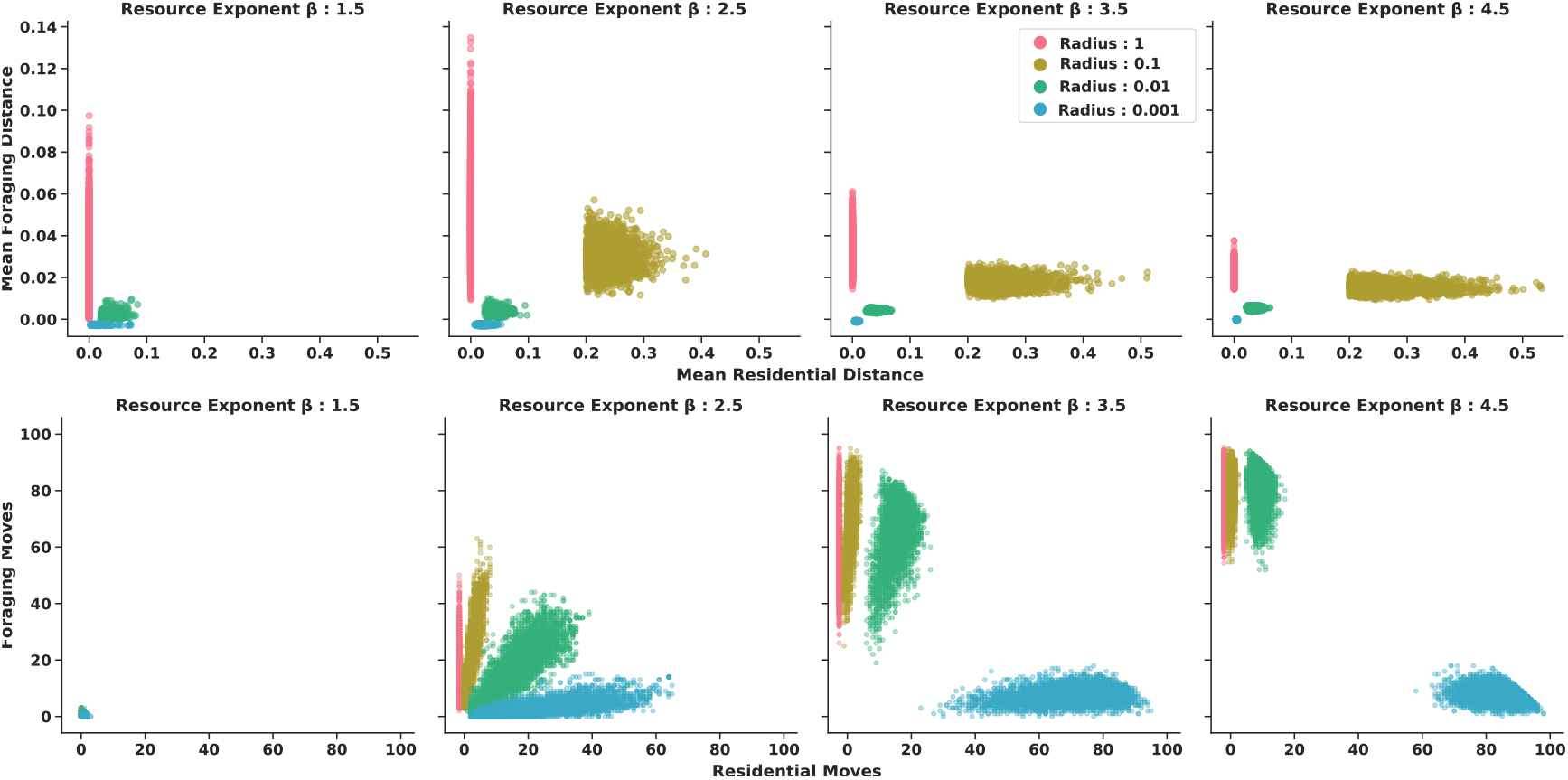
Mobility regimes across different environment and radii. ***Top***: The plot shows the mean distance moved in residential moves (*d*_*r*_) against foraging moves(*d*_*f*_). ***Bottom***: The plot shows the frequency of residential moves(*n*_*r*_) and the frequency of foraging moves(*n*_*f*_).

**Figure 5:**
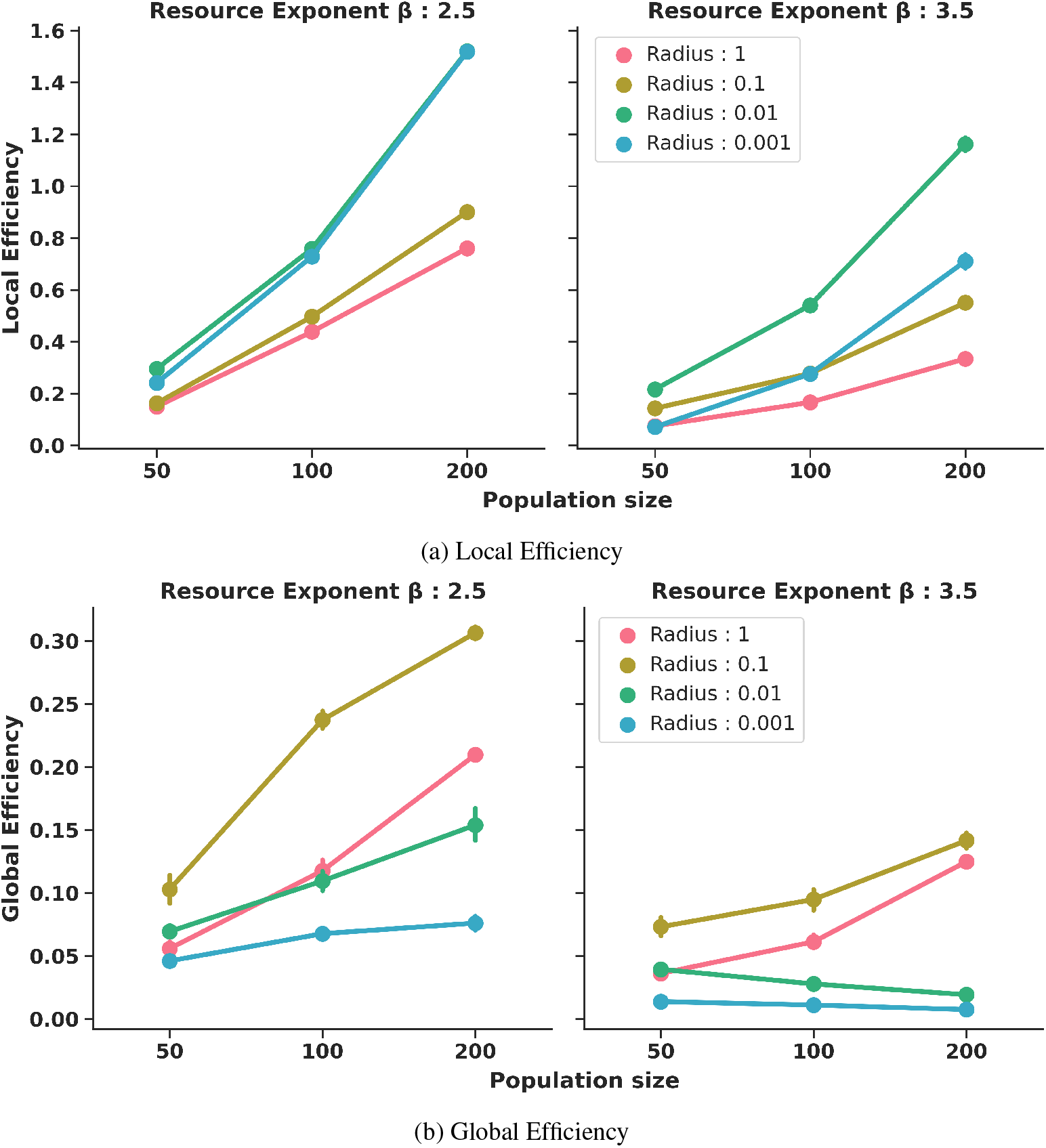
Network efficiencies as a function of population sizes after 100 time-steps for *β* = 2.5 and *β* = 3.5. Error bars indicate 95% confidence intervals.

Moreover, when the environment was more abundant and heterogeneous (*β* = 2.5) and agents could spend longer times on rich patches and formed more locally efficient networks.

On the other hand, we found that the effect of population size on global efficiency was not as straightforward (Fig.5b). In environments with *β* = 2.5, global efficiency increased with an increase in population size. This increase was more exaggerated for central-place foraging (*r >* 0.001) and most substantial for intermediate radius (*r* = 0.1). The longer residential moves enabled more foraging-units to interact and helped create a more connected network. It is also important to note that the networks generated by populations of a small size (*n* = 50) and intermediate foraging-radii were more efficient and connected than networks from large population sizes (*n* = 200) that engaged in ‘point-to-point’ (or *r* ≤ 0.001) foraging.

However, for less abundant and more homogeneous environments (*β* = 3.5), the effect of population size was diminished and only the larger radii (*r* ≥ 0.1) led to an increase in the global efficiencies. When the radii were small, or foraging was similar to ‘point-to-point’, the foraging-units experienced lower encounter rates due to shorter residential moves. Without an increase in long-range connections which would have decreased the path length between network nodes, an increase in population size (or number of network nodes) decreased the global efficiency (Eq. 1) of the networks. When foraging-units moved longer distances and foraged within larger radii, the long-range connections compensated for a larger network and maintained global connectivity even as population sizes increased. Overall, our results suggest an important role of mobility strategies in mediating the effect of population sizes on information transmission.

### 3.4 Central-place foraging networks are efficient at information transmission

**Figure 6:**
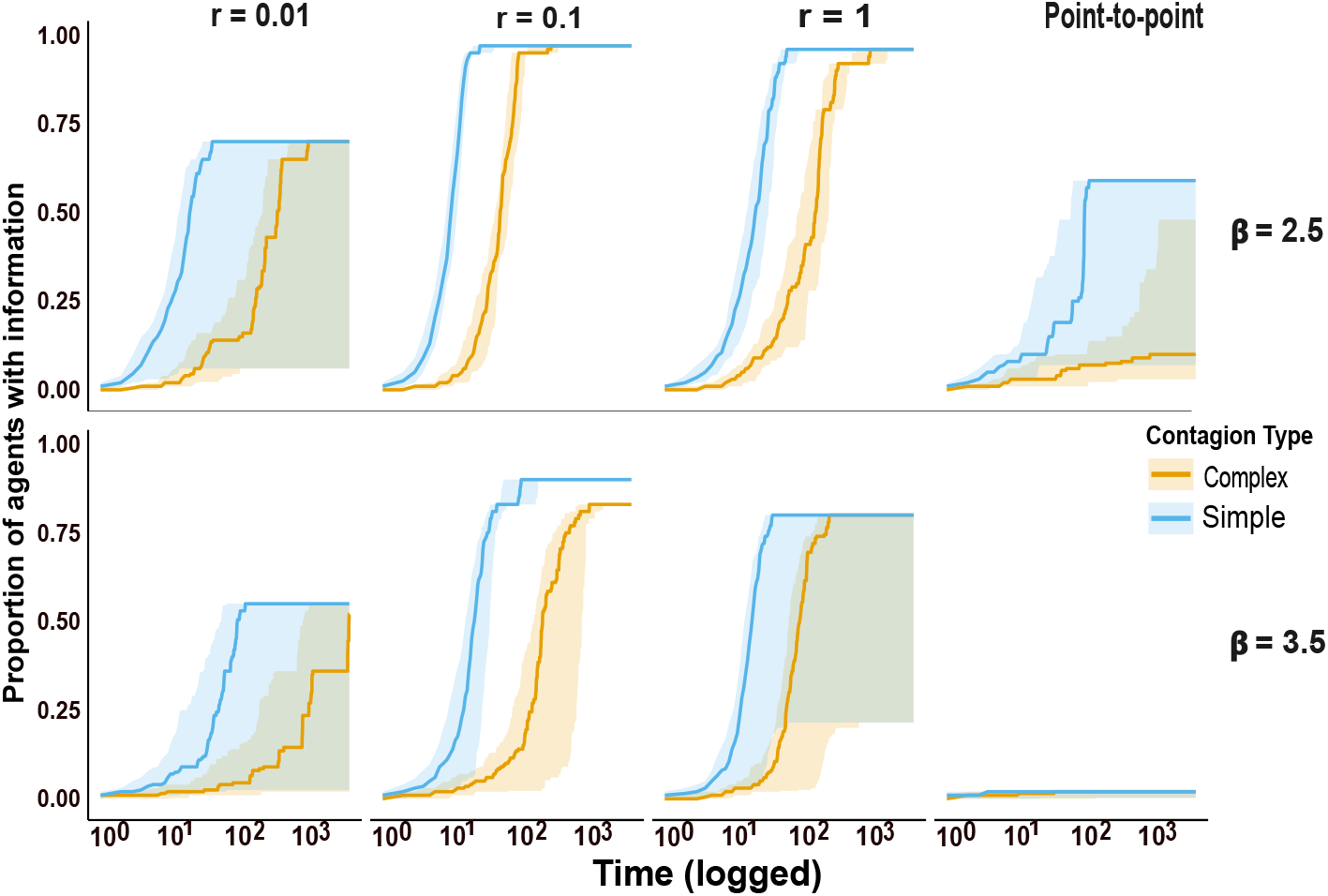
Simple and complex contagion trajectories. The plot shows the spread and speed of contagion over time for different radii (columns) and beta (rows). Shaded regions show the 25^th^ and 75^th^ percentiles of the distribution of trajectories at each time-step.

To directly test the networks for their capability of transmitting information, we conducted both simple and complex contagion simulations on the most globally efficient networks that resulted from each model and parameter combination (45).

In line with efficiency results (see previous section), we found that central-place foraging strategies characterized by a combination of residential and foraging moves (for *r* = 0.1 and *β* = 2.5) formed networks that allowed a rapid diffusion of information, reaching almost every node. We found that information spread more readily in networks with more extensive and well-connected subgroups instead of sparser or fragmented networks. For instance, in the point-to-point model, information reached a maximum of around 50% of the population across environments.

In complex contagion, where multiple novel interactions were required for successful transmission of information, we observed a greater effect of network structures and a slower rate of transmission across networks. For example, for *r* = 0.1 and *β* = 2.5, simple contagion tended to reach 75% of the nodes much faster and more reliably (± SD) 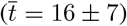 than complex contagion which took longer time to reach similar proportions 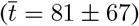 This effect was magnified for less efficient networks (for example, *r* = 1) where the transmission was much slower and more variable 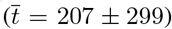 than in simple contagion 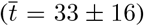 to reach the same proportion (75%) of nodes. In summary, we found that the networks that have high global and local efficiencies (such as those emerging from *β* = 2.5 and *r* = 0.1) can maximize both the reach and speed of contagions that resemble cultural transmission.

## 4 Discussion

Recent work on prehistoric and contemporary hunter-gatherer societies has shown that their social networks are efficient at information transmission and could have accelerated cultural evolution (5; 6; 46). However, the different factors that could have affected the formation of efficient social connectivity are not well understood. In this paper, we focused on assessing how hunter-gatherer foraging patterns could have played a role in the emergence of such efficient social networks. We modeled spatial patterns and mobility regimes emerging from central-place foraging, a derived feature in our lineage, under different environments and tested their implications on the emergence of social networks that are efficient at information transmission (47).

Central-place foraging is characterized by foragers bringing back food to central places (homes) while periodically changing the location of such homes according to the availability of resources. Our results reveal that this foraging pattern could have created social networks that are particularly suited for information exchange. Previous works have suggested that a change in spatial and residence patterns could have caused unique expansions in early hominin social networks (1; 46). We show that, compared to point-to-point foraging, central-place foraging could have modified spatial and residential patterns in ways that would have increased our ancestors’ social interactions, made the networks more expansive and improved their ability to exchange information (22). The main finding from the model by Ramos-Fernandez et al. (35) showed that interactions between ‘point-to-point’ foragers following a basic resource-maximisation rule can result in structured networks with fission-fusion dynamics. Previous works have hypothesized that fission-fusion in non-human primates could have been a precursor for multi-level human social networks (25; 48; 49; 24). Our results show that the addition of central-place foraging can result in a more extensive fission-fusion, larger and more efficient networks, and suggest one possible pathway that could have partly driven such a transition.

We also find support for the idea that environment-driven variability between the mobility regimes employed by different hunter-gatherer societies has significant consequences for their social networks and hence cultural transmission (50; 51; 18; 10). Similar to Perreault and Brattingham (36), we find that mobility regimes which combine short-scale foraging and long-scale residential movements can create more efficient networks as opposed to regimes that are primarily residential or sedentary. In heterogeneous environments, when central-place foragers’ movements are restricted within an intermediate radius with occasional long residential moves to richer resource patches, the networks formed contain densely connected sub-groups embedded in more extensive regional networks. Our results predict that an intermediate mobility regime (Fig 2 inset), thus, could balance the trade-off between networks that are highly cliquish at the expense of global efficiency and sparser large networks that have low clustering. Such networks, similar to small-world topologies, can support information processing at local and global scales (52; 53; 4).

Similar to previous research highlighting the importance of demography for cultural evolution, we find that an increase in population density can result in more efficient networks and a larger capacity for information exchange (54; 55; 56). Our results support previous predictions that population density is not the sole explanation for cultural transmission, and mobility plays an equally important role (56; 57). We show that residential mobility and central-place foraging can improve connectivity even in small populations (58), and they can generate networks that are as efficient as the networks from large populations engaged in ‘point-to-point foraging’. Hunter-gatherer groups with low population densities could have increased their mobility to maintain encounter rates that would have kept them viable by allowing better connectivity, promoting exogamy, efficient exchange of information and resilience to climatic variation (59; 46). Thus, our results also emphasize the importance of optimal connectivity and mobility within a population to offset the adverse effects of demographic changes on cultural transmission (54; 34).

Our results show that ecologically-driven foraging and mobility decisions can generate networks that resemble the structure and composition of networks observed in real-hunter gatherer societies. The agents or foraging units in our model represented nuclear families which across hunter-gatherer societies normally comprise around 4-5 individuals (60; 37). We found that in environments with intermediate heterogeneity (*β* = 2.5), central-place foraging with intermediate radii (*r* = 0.1) that afforded local interactions within overlapping foraging-radii and global interactions due to longer residential moves formed networks with multiple and nested levels. More specifically, the emergent networks fused foraging-units into different (≈ 15) sub-groups (analogous to bands of co-residing family units) that were composed of 5-7 foraging-units each, and on an average half of these sub-groups (7 - 9) were inter-connected with sparse ties forming a higher level of organization of ≈ 40 foraging-units (analogous to communities or mega-bands). Such network organization is similar to ethnographic reports across 336 contemporary hunter-gatherer societies (60; 37) and estimations based on energetic constraints (61; 18; 13; 32; 5) that show hunter-gatherer regional metapopulations of 100 families that can be fragmented into co-residing bands of ≈ 10 families, which are in turn interconnected and form a larger community (≈ 3 − 4 co-residing bands or ≈ 30 families) within the metapopulation. For better comparisons with empirical data on social organization, future studies can base their models on empirical ecological or mobility data and investigate the emergence of multi-level sociality in more detail.

These findings hold significant implications for our species’ evolutionary history and the ability to develop cumulative culture (62). The degree and strength of intra-and inter-regional group interactions among prehistoric hunter-gatherers and their spatial distribution have been proposed to be key factors for cultural transmission (27). The focus of our model was on social network patterns that can arise solely from the derived features of hunter-gatherer foraging-related mobility in different environments, as we wished to unravel their implications for information transmission. Accordingly, we did not consider other social factors that could have shaped their mobility decisions (such as cooperative breeding, resource sharing or joint ritual participation), further structured their interaction networks or potentially resulted in greater incentives and/or efficiency of information transmission. Nonetheless, the model sheds light on the mechanisms by which the regional-scale connectivity generated by individual central-place foraging despite low population sizes throughout our species history. Such connectivity could have maintained cultural diversity and complexity by allowing cultural recombination, transmission of innovations, and preventing the loss of existing culture (11; 63; 2; 5; 64).

Further research could elaborate on more complex portrayals of physical (for example, resource distribution, traveling costs, seasonality) and social (for example, demography, inter-forager competition, cooperation, sociality, kinship, learning) environments that would have characterized early hunter-gatherer communities. These factors would have potentially interacted with foraging and mobility decisions and cultural complexity (65; 66; 67; 68; 29). Moreover, these factors would have also interacted with the cognitive capacities of our early ancestors (e.g. spatial memory, longer-range planning, larger neocortex, theory of mind, symbolic communication) (69; 70). Such cognitive factors would have affected the ability to explore larger spaces, engage in central-place foraging and maintain more extensive social networks, and possibly created selection pressures that paved the way for present-day human cognition and culture (71; 72; 73; 74; 75).

Although additional studies should also address potential selection pressures experienced by our ancestors that would have led them to start using and returning to central places, our study corroborates early claims that central-place foraging would have had important implications for the accumulation and transmission of tools and other types of information (12). However, our work highlights the role of mobility and spatial patterns that stem from central-place foraging in our evolutionary history. We suggest that mobility-driven networks could have led to positive feedback whereby a more efficient transmission of social and/or ecological information, increased food-sharing, better resource-defenses, and a greater accumulation of material culture at a few places would have been advantageous to central-place foragers (76; 39). This advantage could have further promoted reliance on increasingly complex culture and encouraged adaptations to social networks (for example, through kinship or trade) to efficiently generate, transmit and sustain such culture (77; 58; 11; 61; 78; 79).

## Data and code availability

An ODD description of our model and all code required to reproduce the model and results can be found in this Github repository: https://github.com/bleuknight/information_transmission

## Acknowledgments

We thank the Diverse Intelligences Summer Institute (DISI 2020) for inspiring this project, and giving us an opportunity to work together. We also thank anonymous reviewers and editors, Robert Foley, Jacob Foster, Luke Premo, and Paul Smaldino and his lab for their helpful feedback and comments. We are also grateful to Luisa Espinós for help with model visualization.

## Author contributions

**V.B.K**: conceptualization, data curation, formal analysis, funding acquisition, investigation, methodology, software, validation, resources, review and editing; **C.P.I**: conceptualization, data curation, formal analysis, funding acquisition, investigation, methodology, software, visualization, project administration, original manuscript, review and editing; **K.G**: conceptualization, data curation, formal analysis, funding acquisition, investigation, methodology, software, visualization, project administration, resources, validation, original manuscript, review and editing; **N.R.O**: conceptualization, data curation, formal analysis, funding acquisition, methodology, software, visualization, original manuscript, review and editing;

