## Supplementary Information for "Hunter-gatherer foraging networks promote information transmission" (from another version of this preprint: version 1, 2021-04-29)

### Supplementary Material

April 29, 2021

#### 1 Methods

In the original formulation, weights represent distance, meaning information travels more slowly along heavier edges. In our models, weights represent social proximity and, therefore, we use their inverse when calculating efficiency. The underlying logic is the same: networks where all nodes are close to one another are more efficient than those where larger distances must be traveled before connecting any two given vertices.

**Global efficiency**:

Let $N$ be all sets of nodes in the network, $n$ the number of nodes in the network, and $w$ the edge weights.

$$E^w = \frac{1}{n} \sum_{i \in N} \frac{\sum_{j \in N, j \neq i} (d_{ij}^w)^{-1}}{n-1}$$

**Local efficiency**:

Let nodes $i$ and $j$ be connected through edges with weight $w_{ij}$, let $k_i$ be the degree of node $i$, and let $d_{jh}(N_i)$ be the length of the shortest path between $j$ and $h$, including only $i$'s direct ties.

$$E_{loc}^w = \frac{1}{n} \sum_{i \in N} \frac{\sum_{j,h \in N, j \neq i} (w_{ij} w_{ih} [d_{jh}^w(N_i)]^{-1})^{\frac{1}{3}}}{k_i(k_i - 1)}$$

**Louvain community detection**:

Louvain detection method is a hierarchical clustering algorithm, that recursively merges communities into a single node to find clusters with high modularity. It compares the density of connections within a community to the density of connections expected from a random network.

#### 2 Results

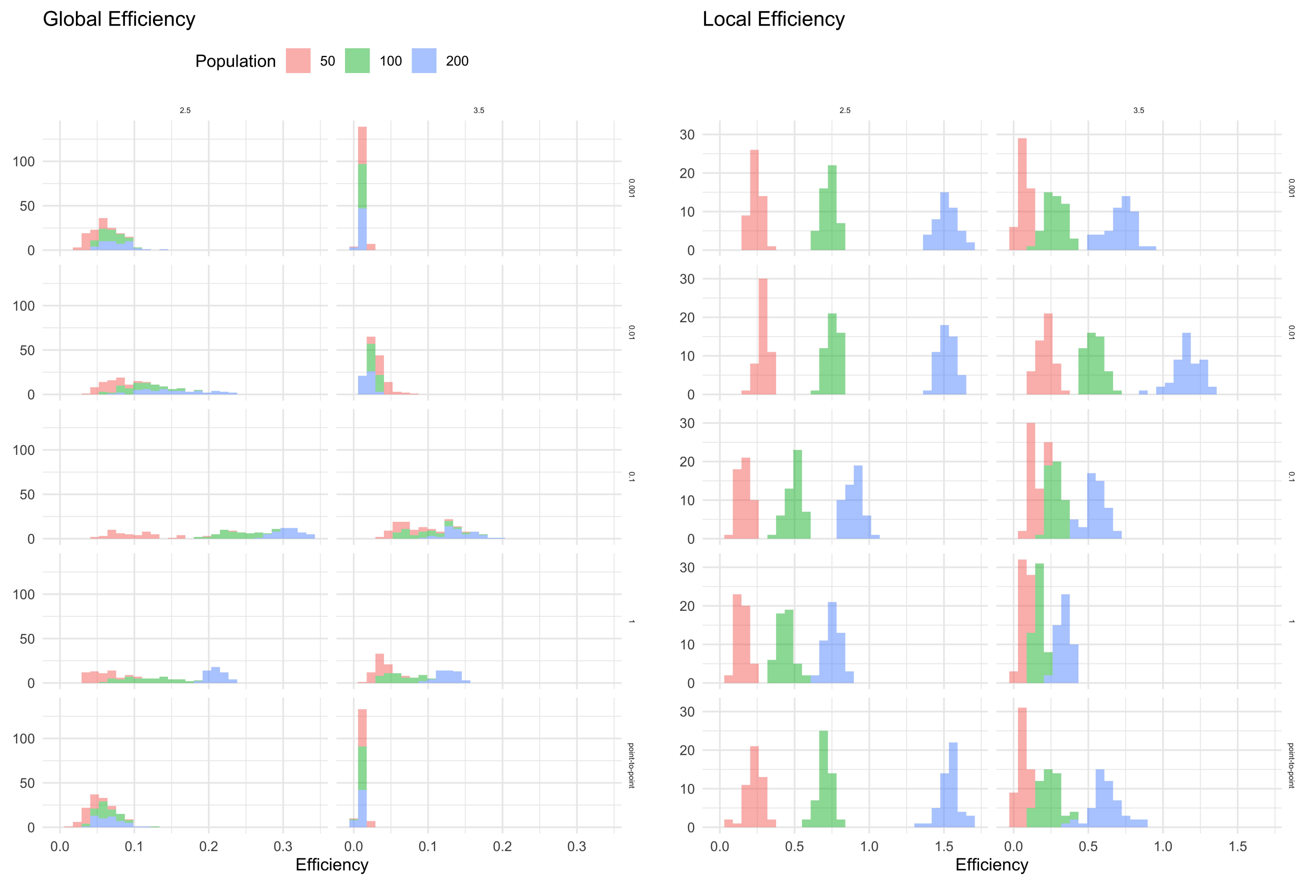


Figure 1: Efficiency distributions from different population sizes (50,100,200 foragers).

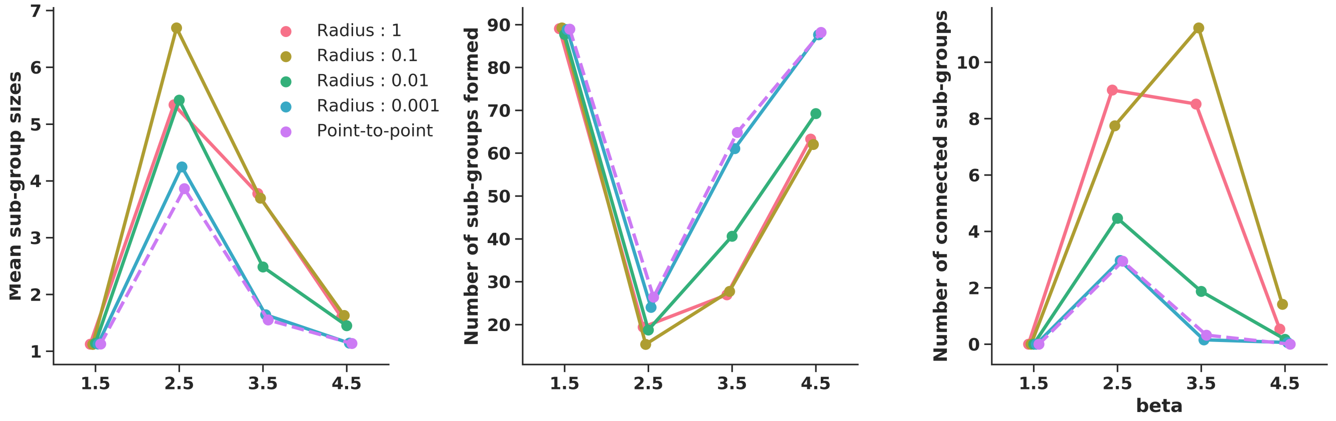


Figure 2: Community structure of networks from 100 foragers.

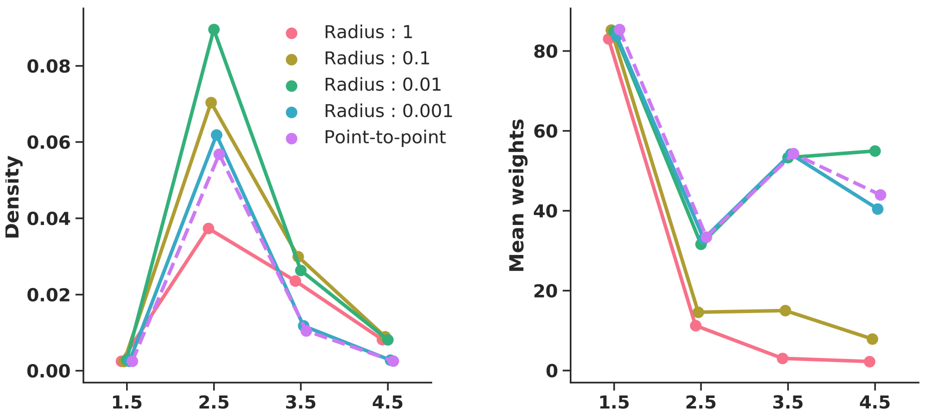


Figure 3: Network features of networks from 100 foragers.

| Beta | Radius | Total number of moves | Total distance moved | Mean distance |
|---|---|---|---|---|
| 1.5 | 0.001 | 1.031±0.304 | 0.02±0.015 | 0.019±0.014 |
| | 0.010 | 1.048±0.319 | 0.027±0.017 | 0.026±0.017 |
| | 0.100 | 1.049±0.306 | 0.02±0.014 | 0.019±0.013 |
| | 1.000 | 1.051±0.329 | 0.021±0.015 | 0.02±0.014 |
| 2.5 | 0.001 | 28.862±14.765 | 0.557±0.295 | 0.02±0.005 |
| | 0.010 | 34.031±15.586 | 0.868±0.422 | 0.026±0.005 |
| | 0.100 | 22.946±12.572 | 1.106±0.706 | 0.046±0.011 |
| | 1.000 | 14.999±7.081 | 0.826±0.634 | 0.05±0.018 |
| 3.5 | 0.001 | 76.814±12.352 | 0.435±0.099 | 0.006±0.001 |
| | 0.010 | 80.359±11.76 | 0.908±0.194 | 0.011±0.002 |
| | 0.100 | 72.522±11.184 | 1.77±0.359 | 0.024±0.003 |
| | 1.000 | 62.169±10.657 | 2.002±0.606 | 0.032±0.007 |
| 4.5 | 0.001 | 90.703±5.534 | 0.357±0.032 | 0.004±0.0 |
| | 0.010 | 90.225±6.536 | 0.732±0.086 | 0.008±0.001 |
| | 0.100 | 85.784±6.282 | 1.568±0.189 | 0.018±0.002 |
| | 1.000 | 84.383±6.486 | 1.723±0.266 | 0.02±0.003 |

Table 1: Summary of Total (foraging and residential) mobility

| Beta | Radius | Total number of F moves | Total F distance | Mean F distance |
|---|---|---|---|---|
| 1.5 | 0.001 | 0.023±0.153 | 0.0±0.0 | 0.001±0.0 |
| | 0.010 | 0.325±0.496 | 0.002±0.003 | 0.005±0.003 |
| | 0.100 | 1.049±0.306 | 0.02±0.014 | 0.02±0.013 |
| | 1.000 | 1.051±0.329 | 0.021±0.015 | 0.02±0.014 |
| 2.5 | 0.001 | 3.277±2.533 | 0.002±0.002 | 0.001±0.0 |
| | 0.010 | 17.703±8.884 | 0.069±0.036 | 0.004±0.001 |
| | 0.100 | 20.925±11.118 | 0.625±0.343 | 0.03±0.007 |
| | 1.000 | 14.999±7.081 | 0.826±0.634 | 0.05±0.018 |
| 3.5 | 0.001 | 7.203±2.712 | 0.005±0.002 | 0.001±0.0 |
| | 0.010 | 64.84±10.021 | 0.271±0.051 | 0.004±0.0 |
| | 0.100 | 70.291±10.623 | 1.205±0.227 | 0.017±0.003 |
| | 1.000 | 62.169±10.657 | 2.002±0.606 | 0.032±0.007 |
| 4.5 | 0.001 | 7.471±2.567 | 0.005±0.002 | 0.001±0.0 |
| | 0.010 | 80.939±6.408 | 0.429±0.056 | 0.005±0.0 |
| | 0.100 | 85.064±6.143 | 1.384±0.23 | 0.016±0.003 |
| | 1.000 | 84.383±6.486 | 1.723±0.266 | 0.02±0.003 |

Table 2: Summary of Foraging mobility

| Beta | Radius | Total number of R moves | Total R distance | Mean R distance |
|---|---|---|---|---|
| 1.5 | 0.001 | 1.008±0.273 | 0.02±0.015 | 0.02±0.014 |
| | 0.010 | 0.723±0.448 | 0.025±0.019 | 0.035±0.012 |
| | 0.100 | 0.0±0.0 | 0.0±0.0 | nan±nan |
| | 1.000 | 0.0±0.0 | 0.0±0.0 | 0.0±0.0 |
| 2.5 | 0.001 | 25.585±12.998 | 0.555±0.294 | 0.022±0.005 |
| | 0.010 | 16.327±7.457 | 0.799±0.394 | 0.048±0.007 |
| | 0.100 | 2.021±1.669 | 0.481±0.402 | 0.237±0.023 |
| | 1.000 | 0.0±0.0 | 0.0±0.0 | 0.0±0.0 |
| 3.5 | 0.001 | 69.611±11.516 | 0.431±0.099 | 0.006±0.001 |
| | 0.010 | 15.52±3.188 | 0.637±0.177 | 0.041±0.006 |
| | 0.100 | 2.232±0.977 | 0.565±0.256 | 0.253±0.033 |
| | 1.000 | 0.0±0.0 | 0.0±0.0 | 0.0±0.0 |
| 4.5 | 0.001 | 83.233±5.736 | 0.352±0.032 | 0.004±0.0 |
| | 0.010 | 9.286±1.735 | 0.303±0.081 | 0.033±0.005 |
| | 0.100 | 0.719±0.605 | 0.184±0.163 | 0.255±0.05 |
| | 1.000 | 0.0±0.0 | 0.0±0.0 | 0.0±0.0 |

Table 3: Summary of Residential mobility

| Beta | Radius | Global Efficiency | Local Efficiency | Network size | Density | Mean Weights |
|---|---|---|---|---|---|---|
| 1.5 | 0.000 | 0.0±0.0 | 0.03±0.03 | 12.17±4.69 | 0.0±0.0 | 85.37±8.5 |
| | 0.001 | 0.0±0.0 | 0.03±0.03 | 12.34±3.42 | 0.0±0.0 | 83.75±7.64 |
| | 0.010 | 0.0±0.0 | 0.05±0.03 | 14.35±3.93 | 0.0±0.0 | 84.87±8.41 |
| | 0.100 | 0.0±0.0 | 0.03±0.03 | 11.99±3.26 | 0.0±0.0 | 85.24±8.69 |
| | 1.000 | 0.0±0.0 | 0.03±0.03 | 12.3±3.82 | 0.0±0.0 | 83.05±8.45 |
| 2.5 | 0.000 | 0.07±0.02 | 0.7±0.04 | 281.23±49.23 | 0.06±0.01 | 33.41±4.28 |
| | 0.001 | 0.07±0.01 | 0.73±0.05 | 306.2±55.64 | 0.06±0.01 | 32.97±3.8 |
| | 0.010 | 0.11±0.03 | 0.76±0.04 | 443.19±106.0 | 0.09±0.02 | 31.62±4.84 |
| | 0.100 | 0.24±0.03 | 0.5±0.05 | 348.35±55.14 | 0.07±0.01 | 14.59±2.01 |
| | 1.000 | 0.12±0.03 | 0.44±0.05 | 184.83±25.96 | 0.04±0.01 | 11.24±0.83 |
| 3.5 | 0.000 | 0.01±0.0 | 0.24±0.07 | 51.78±10.47 | 0.01±0.0 | 54.28±6.46 |
| | 0.001 | 0.01±0.0 | 0.28±0.06 | 58.17±10.07 | 0.01±0.0 | 54.2±6.24 |
| | 0.010 | 0.03±0.01 | 0.54±0.06 | 130.24±20.3 | 0.03±0.0 | 53.29±6.04 |
| | 0.100 | 0.1±0.03 | 0.28±0.04 | 148.2±21.9 | 0.03±0.0 | 14.99±2.21 |
| | 1.000 | 0.06±0.02 | 0.17±0.03 | 116.61±12.87 | 0.02±0.0 | 3.02±0.37 |
| 4.5 | 0.000 | 0.0±0.0 | 0.02±0.02 | 12.55±3.39 | 0.0±0.0 | 43.96±11.59 |
| | 0.001 | 0.0±0.0 | 0.03±0.03 | 13.62±3.58 | 0.0±0.0 | 40.45±11.46 |
| | 0.010 | 0.01±0.0 | 0.17±0.06 | 40.15±7.59 | 0.01±0.0 | 54.96±7.18 |
| | 0.100 | 0.01±0.0 | 0.06±0.04 | 44.17±9.36 | 0.01±0.0 | 7.89±2.74 |
| | 1.000 | 0.01±0.0 | 0.04±0.02 | 40.44±8.07 | 0.01±0.0 | 2.24±0.98 |

Table 4: Summary of network metrics

| Beta | Radius | Number of sub-groups | Mean sub-group size | Number of connected sub-groups |
|---|---|---|---|---|
| 1.5 | 0.000 | 88.93±3.74 | 1.13±0.05 | 0.0±0.0 |
| | 0.001 | 88.87±2.83 | 1.13±0.04 | 0.0±0.0 |
| | 0.010 | 87.71±2.85 | 1.14±0.04 | 0.0±0.0 |
| | 0.100 | 89.23±2.68 | 1.12±0.03 | 0.0±0.0 |
| | 1.000 | 89.08±3.07 | 1.12±0.04 | 0.0±0.0 |
| 2.5 | 0.000 | 26.39±3.59 | 3.86±0.54 | 2.94±1.5 |
| | 0.001 | 24.03±3.42 | 4.25±0.62 | 2.97±1.44 |
| | 0.010 | 18.75±2.41 | 5.42±0.7 | 4.47±1.6 |
| | 0.100 | 15.39±2.61 | 6.69±1.19 | 7.75±1.15 |
| | 1.000 | 19.39±3.35 | 5.34±1.13 | 9.01±1.87 |
| 3.5 | 0.000 | 64.83±4.77 | 1.55±0.12 | 0.32±0.52 |
| | 0.001 | 61.09±3.53 | 1.64±0.1 | 0.16±0.36 |
| | 0.010 | 40.63±3.91 | 2.48±0.24 | 1.87±1.38 |
| | 0.100 | 27.76±4.35 | 3.7±0.61 | 11.22±1.81 |
| | 1.000 | 26.94±3.49 | 3.78±0.51 | 8.52±2.14 |
| 4.5 | 0.000 | 88.18±2.93 | 1.14±0.04 | 0.0±0.0 |
| | 0.001 | 87.64±2.89 | 1.14±0.04 | 0.06±0.24 |
| | 0.010 | 69.22±3.98 | 1.45±0.08 | 0.18±0.38 |
| | 0.100 | 62.04±6.48 | 1.63±0.17 | 1.41±1.25 |
| | 1.000 | 63.28±6.06 | 1.6±0.19 | 0.54±0.96 |

Table 5: Summary of network community structure

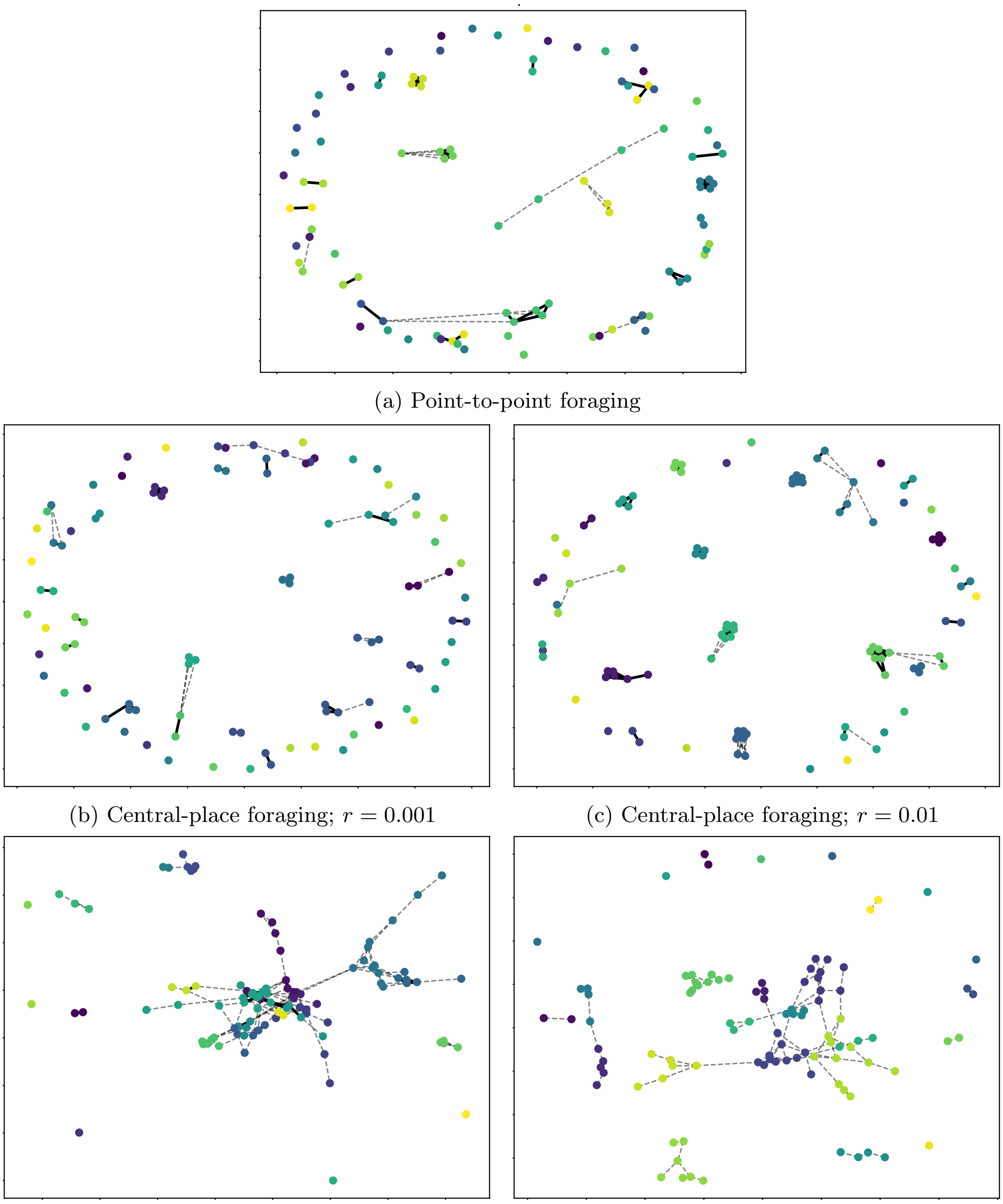

(a) Point-to-point foraging

(b) Central-place foraging; $r = 0.001$

(c) Central-place foraging; $r = 0.01$

(d) Central-place foraging; $r = 0.1$

(e) Central-place foraging; $r = 1$

Figure 4: Emergent Networks from 100 foragers after 100 time-steps. The plot shows an example of weighted networks that emerge from different foraging behaviors in $\beta = 3.5$ environment (clockwise: *Point-to-point*, $r = 0.001, r = 0.01, r = 0.1, r = 1$). Node colors depict the different sub-groups detected by Louvain community-detection method. Edge colors and style depict the edge weights: Solid back edges show strong bonds; dashed grey edges show weak bonds. Distance between nodes also depict the strength of connections. Networks were made using spring layout from *NetworkX* package in Python.
